# INCREASING DIVERSITY AMONG *LENS* SPECIES FOR IMPROVING BIOLOGICAL NITROGEN FIXATION IN LENTIL

**DOI:** 10.1101/2022.10.12.510843

**Authors:** Ana Vargas, Linda Y. Gorim, Kirstin E. Bett

## Abstract

Exotic germplasm is a key resource for reintroducing genetic variability into cultivars. We evaluated 36 accessions from cultivated lentil (*Lens culinaris* Medik.) and six related wild species, inoculated with a commercial strain of *Rhizobium leguminosarum* bv. *viciae* under greenhouse conditions. The objective was to explore *Lens* species and/or accessions that could contribute higher biological nitrogen fixation (BNF) ability to the lentil crop. A split plot design was used with either *Rhizobium* inoculation, added Nitrogen (N) or neither, as the main plots, and accessions in subplots randomized in blocks. Two repeats of the experiment were evaluated at flowering for N fixation and nodulation characters, and two subsequent experiments, with a subset of 14 accessions, were evaluated at maturity for seed production, seed quality and harvest index. Differences in phenotypic expression did not correspond to any particular *Lens* species. CDC Greenstar exhibited some of the highest N fixation values observed among the cultivars and also superior yield results compared to the added N treatment. Wild accessions, including IG 72643 (*L. orientalis*), displayed unique and multiple desirable characteristics compared to cultivars including indeterminate nodulation, higher N translocation, stable yield compared to added N treatment and exceptionally high protein concentration in seeds.

## INTRODUCTION

Wild relatives and related species are the greatest source of genetic diversity for crop breeding, representing a key element for adaptation to changing growing conditions and food security (Global Crop Trust 2019). The introgression of such diversity through interspecific hybridization has resulted in significant progress in the breeding for resistance to biotic and abiotic stresses for the most important grain legume crops such as soybean, peanut, chickpea, lentil and common bean (Muñoz et al., 2017).

Breeding objectives for lentil (*Lens culinaris*) have mainly focused on yield and specific seed characteristics, increasing production but also narrowing genetic variability (Erskine et al., 1998) and reducing the levels of resistance to biotic and abiotic stresses. Differential root traits among genotypes, root distribution and drought tolerance mechanisms have been reported for wild *Lens* accessions (Gorim and Vandenberg 2017). Resistance to diseases such as anthracnose, ascochyta blight and stemphylium blight has been transferred from wild to cultivated lentils through interspecific hybridization (Gela et al., 2021; Vail et al., 2012; Podder et al, 2012; Gupta and Sharma 2006) but linkage drag will also bring in variability in other traits (Ramsay et al., 2021) – some positive, but not all.

The biological nitrogen fixation process (BNF) contributes about a third of the nitrogen (N) that is used in agroecosystems annually (Herridge et al., 2008). This is converted into protein available for human consumption as fresh vegetables and dried grains but also leaves below-ground contributions to subsequent crops (Aslam et al., 2003). The proportion of N derived from the atmosphere (Ndfa) in lentil averages around 65%, but a broad range of estimates have been reported from 9-97%, with values of 4 to 152 Kg ha^−1^ of shoot N (Peoples et al., 2009).

Modern legume varieties are usually selected under high fertility conditions, where all necessary nutrition is provided in available forms from synthetic fertilizers, making it unnecessary to establish symbiotic relations, potentially contributing to the loss of genes that control these traits (Wissuwa et al., 2009). In common bean (*Phaseolus vulgaris*), the contributions from crosses with the related tepary bean (*Phaseolus acutifolius*) through interspecific hybridization have resulted in germplasm with higher capacity to fix N (Somasegaran and Hoben 1991). In contrast, in soybean (*Glycine max*), higher N fixing ability may have been gained during the domestication and breeding processes, based on a study of a group of wild and cultivated genotypes, as well as a recombinant inbred line population (Muñoz et al., 2016).

Improving the legume -rhizobia symbiosis is the most important route for increasing efficient use of N in cropping systems. Diverse lentil cultivars have been tested under controlled conditions with *Rhizobium leguminosarum* bv. *viciae*, with significant variation noted for the proportion of Ndfa, suggesting the possibility of breeding for increasing N fixation (Abi-Ghanem et al., 2011). Contributions may also be made to the N fixing ability of lentil from its wild relatives. This study was designed to identify species and genotypes from a broad set of accessions from the genus *Lens* that could be useful in breeding for better BNF ability.

## MATERIALS AND METHODS

### Accessions and *Rhizobium* strain

The 36 accessions that were tested, representing a cross section of all *Lens* species, are listed in Table 1. The inoculant used for the experiment was the strain BASF 1435 Nodulator XL^®^ *Rhizobium leguminosarum* bv. *viciae* (*Rlv*), a commercial inoculum currently used by farmers. This strain would constitute part of the resident population of rhizobia in pulse-growing areas of the Northern Great Plains as it has been used for at least two decades.

**Table 1.** Accession, species, gene pool and origin of 36 *Lens* accessions studied for their N fixing ability following inoculation with BASF 1435 Nodulator XL^®^ (*Rhizobium leguminosarim* bv. *<u>viciae</u>*<u>) using the difference method.</u>

| Accession | Species | Gene Pool | Country of Origin/Source |
| --- | --- | --- | --- |
| <u>Cultivated</u> |  |  |  |
| CDC Maxim<sup>a</sup> | *L. culinaris* | Primary | Canada |
| CDC Redberry | *L. culinaris* | Primary | Canada |
| CDC Robin | *L. culinaris* | Primary | Canada |
| CDC Milestone | *L. culinaris* | Primary | Canada |
| CDC Asterix | *L. culinaris* | Primary | Canada |
| CDC KR-1<sup>a</sup> | *L. culinaris* | Primary | Canada |
| CDC Greenstar<sup>a</sup> | *L. culinaris* | Primary | Canada |
| CDC QG-4 | *L. culinaris* | Primary | Canada |
| Eston | *L. culinaris* | Primary | Canada |
| VIR-421 | *L. culinaris* | Primary | Russia |
| Lupa<sup>a</sup> | *L. culinaris* | Primary | Spain |
| ILL 7502 | *L. culinaris* | Primary | ICARDA<sup>b</sup> |
| ILL 1704 | *L. culinaris* | Primary | ICARDA |
| ILL 8006 | *L. culinaris* | Primary | ICARDA |
| Indianhead<sup>a</sup> | *L. culinaris* | Primary | Canada |
| <u>Wilds</u> |  |  |  |
| BGE 016880<sup>a</sup> | *L. orientalis* | Primary | Israel |
| IG 72529 | *L. orientalis* | Primary | Turkey |
| IG 72611 | *L. orientalis* | Primary | Turkey |
| IG 72622 | *L. orientalis* | Primary | Turkey |
| IG 72643<sup>a</sup> | *L. orientalis* | Primary | Syria |
| IG 72672 | *L. orientalis* | Primary | Turkey |
| PI 572376<sup>a</sup> | *L. orientalis* | Secondary | Turkey |
| IG 72613 | *L. tomentosus* | Secondary | Turkey |
| IG 72614 | *L. tomentosus* | Secondary | Turkey |
| IG 72805 | *L. tomentosus* | Secondary | Turkey |
| PI 572390<sup>a</sup> | *L. tomentosus* | Secondary | Turkey |
| IG 72543 | *L. odemensis* | Secondary | Palestine |
| IG 72623<sup>a</sup> | *L. odemensis* | Secondary | Turkey |
| IG 72760<sup>a</sup> | *L. odemensis* | Secondary | Syria |
| IG 110810<sup>a</sup> | *L. lamottei* | Secondary | Spain |
| IG 110813 | *L. lamottei* | Secondary | Spain |
| IG 72537 | *L. lamottei* | Secondary | France |
| IG 72815 | *L. ervoides* | Tertiary | Turkey |
| L01-827A<sup>a</sup> | *L. ervoides* | Tertiary | Unknown<sup>d</sup> |
| LR59-81<sup>ac</sup> | *L. culinaris* × *L. ervoides* | interspecific | Canada<sup>d</sup> |
| IG 116024 | *L. nigricans* | Quaternary | Turkey |

<sup>a</sup>evaluated at maturity, <sup>b</sup>ICARDA: International Center for Agricultural Research in the Dry Areas, <sup>c</sup>LR59-81: interspecific line (Eston × L01-827A), CDC, University of Saskatchewan, Canada. <sup>d</sup>Fiala et al., 2009.

### Growing conditions and inoculum

The experiments were conducted in the greenhouse facilities at the University of Saskatchewan (USask). The chamber was set to a 16-hour day length and temperature was set to 25 □C during the day and 18 □C at night. Two repeats of the experiment were planted for evaluation at flowering and an additional two repeats, with a subset of 14 accessions (indicated in Table 1), were planted for evaluation at maturity. About 30 seeds per accession were disinfected, scarified and pre-germinated in soft agar (6% w/v) 48-72 hours before planting. For disinfection, seeds were surface sterilized with 70% ethanol (v/v) for 30 seconds, followed by 5% bleach (v/v) for 2 minutes, and washed with running distilled water. Scarification was conducted to ensure germination of the wild accessions and was carried out by nicking the seed coat with a razor blade before plating. A low-N growing mix, composed of Sunshine^®^ #3 (Sungro Horticulture, Agawan, MA) and sand (1:1 v/v), pasteurized for 72 hours at 70 □C, was used to ensure the absence of rhizobia and other microbial populations. N-free nutrient solution (Somasegaran and Hoben 1994) was applied to the growing media in polyvinyl chloride (PVC) cylinders (11 cm diameter × 40 cm depth) 24 hours before the germinated seeds were planted. The chemical N treatment consisted of a split application of Co (NH_2_)_2_ (Urea), applying 25% along with the nutrient solution and 75% two weeks later (219 mg of N per cylinder total).

^a^evaluated at maturity, ^b^ICARDA: International Center for Agricultural Research in the Dry Areas, ^c^LR59-81: interspecific line (Eston × L01-827A), CDC, University of Saskatchewan, Canada. ^d^Fiala et al., 2009.

All treatments had optimal fertility for all the other elements to ensure full N fixation potential. Amounts of nutrients per cylinder were: 529 mg of P, 402 mg of K, 53 mg of Mg, 49.5 mg of Zn, 53 mg of Ca, 1.95 mg of Mo, 1.85 mg Mn, 2.15 mg of Cu, 6.7 mg Fe, and 1.8 mg of Co. Sources of fertilizers were: KH_2_PO_4_, triple superphosphate (TSP), MgSO_4_.7H_2_O, CaCl_2_, ZnSO_4_.7H_2_0, Mo_7_O_24_.H_2_O, MnSO_4_.H_2_O, CuSO_4_.5H_2_O, H_3_BO_3,_ FeC_6_H_5_O_7_.3H_2_O and CoSO_4_.7H_2_O. The same amount of water was applied to each cylinder: from 0-7 DAS 200 ml total, from 8-14 days 70 ml daily, from 14 until flowering 150 ml daily. For the experiments evaluated until maturity, 250 ml of water were added daily from flowering to maturity.

For the rhizobia treatment (R), pure *Rhizobium* culture grown on yeast mannitol agar (Sigma Aldrich, St. Louis, MO) was transferred to yeast mannitol broth (Sigma Aldrich, St. Louis, MO). The liquid culture was agitated in a shaker (Orbital 420, Thermo Scientific, Waltham, MA) for 48 hours at 26 □C and 180 rpm. Liquid inoculum was standardized to 3×10^9^ cells/ml and inoculation was conducted 24 hours after transferring the germinated seeds to the growing medium by applying 1 ml of liquid inoculum to the growing media of each cylinder.

### Experimental Design and Statistical Analysis

Treatments were arranged in a split-plot design to prevent infection of rhizobia in the Control (C) and added N (+N) treatments, with each cylinder representing an experimental unit. Treatments were the main plots: R: *Rhizobium*, no added N; Control: no *Rhizobium*, no added N; +N: added N, no *Rhizobium*. The 36 lentil accessions were randomized to the subplots in four blocks, for a total of 432 experimental units and the whole experiment was repeated twice. The same treatments, distribution and replicates were used for two additional repeats evaluated at maturity, with 14 selected accessions (Table 1), for a total of 168 experimental units.

An ANOVA was conducted with main plots and accessions considered as fixed and blocks and repetitions as random to test for significance of effects. Means were separated with the least significant difference test (LSD; P_≤_0.05) using the statistical package SAS (SAS Institute, 2015). A Pearson correlation matrix was also generated among all measured parameters in SAS. It was conducted with the 14 accessions evaluated at maturity to establish relationships among parameters evaluated at flowering and maturity. Results from the C plot were used only for the parameters related to N fixed.

### Phenotyping

Because of the variation in phenological cycles, plants were evaluated as they flowered (first open flower), to ensure they were all the same stage when the measurements were taken. Days to flowering (DTF) were recorded. Shoots were separated from roots with scissors and dried at 70 □C for 72 hours to determine shoot dry weight (SDW). A shoot subsample was ground and passed through a 2 mm mesh screen and N concentration in shoots was determined on 100 mg samples using a LECO analyzer (Leco FP628, Leco Corporation, St. Joseph, MI, USA). N accumulation was estimated as a product of N concentration in shoots and SDW. These values were used to determine the N fixed by using the N difference method as N_2_ fixed = [N yield inoculated plant (N/100*SDW) - N yield control plant (N/100*SDW)] (Unkovich *et al*., 2008). Each accession was included in the Control and N fixation was estimated using the matching genotype as the reference plant. Roots of plants in the Control were examined to ensure absence of nodulation. Roots were washed using a 5 mm screen separating all growing media while keeping the entire root system. Washed roots were placed in plastic bags in a fridge at 6 □C to prevent dehydration prior to evaluation. Presence of nodulation was scored by counting the nodule number (NN), and nodule dry weight (NDW) was estimated following desiccation at 60 □C for 48 hours. After harvesting the nodules, roots were scanned (Epson Perfection V 700, Regent Instruments Inc., QC, Canada) and analyzed using WinRhizo (Regent Instruments Inc., Quebec, QC, Canada) to determine total root length (TRL), root average diameter (RAD), total root surface area (TRSA), and total root volume (TRV). Roots were then dried at 70 □C for 72 hours and root dry weight (RDW) was recorded. Specific nodule weight (SpNDW) was estimated by dividing NDW by NN as a measure of nodule size, and root to shoot ratio (R:S) was calculated using RDW and SDW. N-use efficiency was calculated in terms of N_2_ fixed/grams of nodule weight. IG 116024 (*L. nigricans*) did not flower before the experiment was terminated; therefore, phenotypic evaluations were conducted 62 days after seeding (DAS) for this accession.

For the subset of 14 accessions taken to maturity, days to maturity (DTM) were recorded for each unit when about 90% of the pods were brown. Whole plants were harvested and dried at 50 □C for 72 hours. Seeds were separated, counted (SN) and weighed to determine seed weight (SW), and the rest of the plant was weighed to obtain plant dry weight. Harvest index (HI) was estimated as (SW/whole plant dry weight) × 100. Seed sub-samples (approx. 30-50 seeds each) were ground to 2 mm with a seed grinder (Cyclone Sample Mill, Seedburo Co., Chicago, IL, USA) and seed protein percentage was determined on 100 mg samples using a LECO analyzer (LECO FP628, Leco Corporation, Saint Joseph, MI, USA). Thousand seed weight (KSW) was estimated as (SW/SN) × 1000.

## RESULTS

### Nodulation and N fixation at flowering

All accessions produced nodules when inoculated with the commercial *Rhizobium* (Table 2). Most cultivated accessions had significantly higher numbers of nodules (NN), ranging from 23 to 451 nodules per plant (See Supplemental Table S1 for ANOVA), with an average of 233.1. Wild accessions had lower NN values, ranging from 5 to 172 per plant, with an average 51.9. Lupa and ILL 7502 were the only cultivated accessions with low NN values, similar to those found in the wilds. Among the wild lentil genotypes, only IG 72623 (*L. odemensis*) and IG 72760 (*L. odemensis*) had NN values similar to some of the cultivated accessions.

**Table 2.** Number of nodules, total nodule dry weight (g), specific nodule weight (mg) and N_2_ fixed (mg/plant) for 36 *Lens* accessions inoculated with the strain BASF 1435 Nodulator XL^®^ (*Rhizobium leguminosarum* bv. *viciae*), calculated using the N difference method (mean±SD).

| Accession | Species | Number of nodules | Nodule dry weight (mg) | Specific nodule dry weight (mg) | N<sub>2</sub> fixed<sup>a</sup> (mg/plant) |
| --- | --- | --- | --- | --- | --- |
| <u>Cultivated</u> |  |  |  |  |  |
| CDC Maxim | *L. culinaris* | 451 (±191) | 85 (±37.1) | 0.19 (±0.05) | 62.3 (±30.8) |
| CDC Redberry | *L. culinaris* | 436 (±139.6) | 129.1 (±54.9) | 0.29 (±0.04) | 72.4 (±23.4) |
| CDC Robin | *L. culinaris* | 192 (±86.4) | 58.0 (±27.8) | 0.30 (±0.08) | 77.2 (±18.3) |
| CDC Milestone | *L. culinaris* | 129 (±13.4) | 33.4 (±12.4) | 0.25 (±0.07) | 76.3 (±30.6) |
| CDC Asterix | *L. culinaris* | 189 (±57.5) | 49.7 (±16.8) | 0.28 (±0.11) | 86.6 (±32.2) |
| CDC KR-1 | *L. culinaris* | 364 (±96.7) | 77.5 (±31.8) | 0.20 (±0.05) | 93.1 (±31.8) |
| CDC Greenstar | *L. culinaris* | 301 (±90.3) | 104.4 (±49.6) | 0.32 (±0.09) | 91.5 (±38.9) |
| CDC QG-4 | *L. culinaris* | 216 (±143.8) | 38.9 (±28) | 0.13 (±0.04) | 50.6 (±16.7) |
| Eston | *L. culinaris* | 206 (±53) | 28.8 (±12.8) | 0.13 (±0.04) | 35.6 (±6.7) |
| VIR-421 | *L. culinaris* | 168 (±47.5) | 27.5 (±11.7) | 0.16 (±0.03) | 26.0 (±15.5) |
| Lupa | *L. culinaris* | 23 (±12.0) | 1.8 (±0.6) | 0.07 (±0.01) | 3.6 (±1.8) |
| ILL 7502 | *L. culinaris* | 99 (±59.8) | 17.7 (±3.7) | 0.25 (±0.12) | 29.0 (±12.1) |
| ILL 1704 | *L. culinaris* | 153 (±64.1) | 22.6 (±3.8) | 0.17 (±0.05) | 19.5 (±9.1) |
| ILL 8006 | *L. culinaris* | 268 (±89.4) | 58.7 (±14.7) | 0.23 (±0.04) | 33.0 (±13.9) |
| Indianhead | *L. culinaris* | 294 (±76.9) | 60.8 (±11.3) | 0.23 (±0.09) | 82.0 (±30.4) |
| <u>Wilds</u> |  |  |  |  |  |
| BGE 016880 | *L. orientalis* | 11 (±8.9) | 16.8 (±2.4) | 0.25 (±0.05) | 5.5 (±7.6) |
| IG 72529 | *L. orientalis* | 19 (±2.9) | 3.5 (±0.8) | 0.20 (±0.09) | 1.5 (±0.43) |
| IG 72611 | *L. orientalis* | 74 (±17.8) | 17.4 (±10.6) | 0.18 (±0.08) | 7.9 (±3.1) |
| IG 72622 | *L. orientalis* | 36 (±12.2) | 11.6 (±4.8) | 0.38 (±0.11) | 4.5 (±2.8) |
| IG 72643 | *L. orientalis* | 39 (±15.2) | 12.4 (±7.4) | 0.33 (±0.11) | 18.6 (±7.1) |
| IG 72672 | *L. orientalis* | 18 (±8.2) | 5.7 (±3.44) | 0.43 (±0.13) | 0.6 (±0.29) |
| PI 572376 | *L. orientalis* | 5 (±1.5) | 4.1 (±1.6) | 0.94 (±0.3) | 3.5 (±2.7) |
| IG 72613 | *L. tomentosus* | 46 (±19.1) | 10.5 (±4.0) | 0.23 (±0.02) | 4.7 (±2.4) |
| IG 72614 | *L. tomentosus* | 19 (±6.8) | 4.28 (±1.63) | 0.27 (±0.12) | 0.6 (±0.2) |
| IG 72805 | *L. tomentosus* | 18 (±7.6) | 6.7 (±1.9) | 0.43 (±0.13) | 7.1 (±3.2) |
| PI 572390 | *L. tomentosus* | 39 (±4.6) | 8.18 (±1.2) | 0.21 (±0.01) | 11.0 (±6.1) |
| IG 72543 | *L. odemensis* | 45 (±14.2) | 9.28 (±3.3) | 0.20 (±0.02) | 6.3 (±2.7) |
| IG 72623 | *L. odemensis* | 143 (±76.2) | 20.5 (±11.1) | 0.14 (±0.03) | 12.4 (±6.2) |
| IG 72760 | *L. odemensis* | 172 (±12.7) | 31 (±8.1) | 0.18 (±0.04) | 35.2 (±15.6) |
| IG 110810 | *L. lamottei* | 17 (±5.7) | 7.18 (±1.7) | 0.46 (±0.09) | 6.1 (±1.9) |
| IG 110813 | *L. lamottei* | 18 (±3.8) | 5.1 (±3.2) | 0.3 (±0.11) | 4.7 (±2.3) |
| IG 72537 | *L. lamottei* | 12 (±3.2) | 4.9 (±2.6) | 0.46 (±0.19) | 1.5 (±0.4) |
| IG 72815 | *L. ervoides* | 67 (±27.1) | 7.23 (±3.6) | 0.11 (±0.02) | 3.4 (±1.4) |
| L01-827A | *L. ervoides* | 13 (±6.4) | 8.05 (±3.7) | 0.7 (±0.21) | 3.3 (±1.3) |
| LR59-81 | interspecific<sup>b</sup> | 127 (±81.2) | 19.4 (±9.4) | 0.1 (±0.01) | 10.6 (±6.3) |
| IG 116024 | *L. nigricans* | 78 (±15.8) | 25.6 (±6.1) | 0.3 (±0.12) | 9.9 (±5.8) |
| LSD |  | 114.2*** | 29.72*** | 0.28*** | 25.56*** |

<sup>b</sup>*L. culinaris* × *L. ervoides*.

Nodule dry weight (NDW) on a whole plant basis was significantly different among accessions (Table 2) with the cultivated ones generally, but not always, having higher NDW than the wilds. NDW was not significantly different among the wild accessions, however. Specific nodule dry weight (SpNDW) was highest in a few wild accessions: PI 572376 (*L. orientalis*), L01-827A (*L. ervoides*), IG 72537 and IG 110810 (*L. lamottei*). The efficiency of nodulation (N fixed per g of nodule weight), however, was not significantly different among accessions.

N fixed per plant (mg) had a great range of values (Table 2) especially for the cultivars: from 1.8-129.13 mg/plant and 54.4 mg/plant on average. Among the wild accessions, values ranged from 0.6-35.2 mg/plant and 8.6 mg/plant on average.

ANOVA values are ns: not significant, ^*, **, ***^: significant at 0.05, 0.01, 0.001 respectively. ^a^N_2_ fixed estimated by the difference method for R plot (Total N accumulation R plot - Total N accumulation C plot). ^b^*L. culinaris × L. ervoides*.

### Effects of inoculation on seed traits and harvest index

A significant effect of *Rhizobium* (R) inoculation compared to the added Nitrogen (+N) treatment, and differences among accessions, as well as some specific accession *×* treatment interactions, were observed for all seed parameters evaluated (Table 3, see Supplemental Table S1 for ANOVA).

**Table 3.** Seed number, seed weight (g), seed protein percentage, thousand seed weight (g), and harvest index of 14 *Lens* accessions inoculated with the strain BASF 1435 Nodulator XL^®^ (*Rhizobium leguminosarum* bv. *viciae*) or added Nitrogen treatments (mean ± SD).

| Accession | Seed number |  | Seed weight (g) |  | Seed protein percentage |  | Thousand seed weight (g) |  | Harvest index (%) |  |
| --- | --- | --- | --- | --- | --- | --- | --- | --- | --- | --- |
|  | +Nitrogen | *Rhizobium* | +Nitrogen | *Rhizobium* | +Nitrogen | *Rhizobium* | +Nitrogen | *Rhizobium* | +Nitrogen | *Rhizobium* |
| <u>Cultivated</u> |  |  |  |  |  |  |  |  |  |  |
| CDC Maxim | 217.7 (±25.9) | 124.3 (±34.1) | 8.5 (±0.8) | 5.3 (±1.1) | 19.1 (±0.8) | 17.0 (±0.7) | 39.1 (±1.9) | 42.3 (±2.9) | 50 (±2.6) | 47 (±4.9) |
| CDC KR-1 | 188.7 (±34.2) | 136.5 (±37.1) | 9.7 (±1.2) | 6.8 (±1.5) | 18.9 (±0.8) | 18.5 (±0.8) | 52.0 (±8.1) | 50.6 (±6.3) | 53 (±4.6) | 56 (±3.1) |
| CDC Greenstar | 139.7 (±8.8) | 125.0 (±18.9) | 9.8 (±0.7) | 9.4 (±1.6) | 19.4 (±0.7) | 18.1 (±0.8) | 70.0 (±4.4) | **75.8** (±6.6) | 55 (±4.8) | 53 (±6.8) |
| Lupa | 103.5 (±22.3) | 43.0 (±9.5) | 3.5 (±1.1) | **1.6** (±0.6) | 19.6 (±0.7) | 17.9 (±0.6) | 33.1 (±3.7) | 35.6 (±3.1) | 43 (±10.5) | 40 (±7.8) |
| Indianhead | 382.5 (±72.9) | 371.2 (±60.2) | 9.1 (±1.9) | 9.8 (±1.4) | 22.6 (±0.9) | 25.3 (±1.0) | 23.8 (±1.1) | 26.6 (±2.7) | 52 (±4.9) | 49 (±2.6) |
| <u>Wilds</u> |  |  |  |  |  |  |  |  |  |  |
| BGE 016880 | 208.5 (±47.5) | 105.3 (±23.4) | 2.9 (±0.8) | 1.8 (±0.4) | 20.3 (±0.8) | 23.8 (±0.9) | 13.8 (±1.6) | 16.6 (±1.0) | 37 (±7.1) | 36 (±1.9) |
| IG 72643 | 279.7 (±78.1) | 207.7 (±42.3) | 3.9 (±1.4) | 2.9 (±0.7) | 21.8 (±0.8) | **26.4** (±1.0) | 14.4 (±3.2) | 14.0 (±2.2) | 35 (±7.9) | 45 (±1.6) |
| PI 572376 | **372.2** (±43.4) | 169.6 (±36.2) | 2.7 (±1.3) | 1.3 (±0.6) | 22.5 (±0.9) | 21.5 (±0.9) | 7.4 (±0.6) | 7.9 (±0.4) | 34 (±5.2) | 37 (±6.8) |
| PI 572390 | 226.0 (±51.2) | 236.3 (±48.5) | 2.9 (±0.6) | 3.2 (±0.8) | 20.8 (±0.8) | **26.2** (±1.0) | 13.0 (±1.6) | 13.8 (±1.2) | 37 (±8.2) | **53** (±7.8) |
| IG 72623 | 357.0 (±71.9) | 329.0 (±70.9) | 5.5 (±1.8) | 5.0 (±1.6) | 24.9 (±1.0) | 24.5 (±1.0) | 14.8 (±1.4) | 15.6 (±1.1) | 40 (±7.7) | 48 (±2.4) |
| IG 72760 | 178.5 (±80.2) | 141.3 (±56.4) | 2.2 (±0.9) | 1.7 (±0.7) | 19.4 (±0.8) | 19.2 (±0.8) | 12.1 (±0.6) | 12.0 (±0.3) | 43 (±10.9) | 46 (±8.9) |
| IG 110810 | 174.8 (±65.0) | 123.2 (±25.8) | 2.6 (±0.8) | 2.2 (±0.9) | 26.2 (±1.1) | 25.8 (±1.0) | 15.1 (±1.0) | 17.2 (±3.9) | 35 (±3.7) | **49** (±9.3) |
| L01-827A | 303.7 (±57.3) | 186.8 (±17.9) | 2.1 (±0.4) | 1.2 (±0.1) | 22.1 (±0.8) | 24.1 (±0.9) | 6.9 (±0.2) | 6.5 (±0.3) | 43 (±1.7) | 44 (±2.0) |
| LR59-81 | 176.5 (±73.7) | 244.7 (±50.2) | 2.6 (±1.2) | 2.3 (±0.8) | 25.7 (±1.1) | 26.5 (±1.1) | 14.4 (±2.0) | 15.5 (±0.7) | 26 (±5.4) | 27 (±4.2) |
| Mean | 236.4 | 174.6 | 4.9 | 3.8 | 25.7 | 26.5 | 23.6 | 25.0 | 42 | 45 |
| LSD | 81.3*** |  | 1.3*** |  | 0.7*** |  | 4.4*** |  | 7.6*** |  |

ANOVA values are ns: not significant, \*, \*\*, \*\*\*: significant at 0.05, 0.01, 0.001 respectively. **Bolded** numbers represent the significant interactions for accession × treatment (+Nitrogen/*Rhizobium*).

Seed number (SN) was lower when inoculated (R) compared to the +N treatment for only a few accessions: CDC Maxim, BGE 016880 (*L. orientalis*) and L01-827A (*L. ervoides*). PI 572376 (*L. orientalis*) had more than twice the SN under +N compared to R, and was the only wild accession with a lower seed weight (SW) under R. All other wilds had SW values that were similar under both treatments. Two cultivated accessions had lower SW when inoculated (CDC Maxim and CDC KR-1) while the rest were no different to the +N treatment results.

CDC Greenstar was the only accession that had higher thousand seed weight (KSW) value under R treatment compared to +N (Table 3). Inoculation had no impact on harvest index (HI) among the cultivated accessions, but IG 72643 (*L. orientalis*), PI 572390 (*L. tomentosus*), IG 72623 (*L. odemensis*) and IG 110810 (*L. lamottei*) had significantly increased HI values under the R treatment.

Seed protein percentage was the seed trait with the most variation observed under R treatment when compared to plants under +N treatment (Table 3). Indianhead, BGE 016880 (*L. orientalis*), PI 572390 (*L. tomentosus*), IG 72643 (*L. orientalis*), IG 72623 (*L. odemensis*), L01-827A (*L. ervoides*) and LR59-81 (interspecific) had increased values when inoculated. IG 72643 (*L. orientalis*) and PI 572390 (*L. tomentosus*) had the biggest seed protein percentage increases when inoculated compared to +N. In contrast, CDC Maxim, CDC Greenstar and Lupa had lower values when inoculated.

ANOVA values are ns: not significant, ^*, **, ***^: significant at 0.05, 0.01, 0.001 respectively. **Bolded** numbers represent the significant interactions for accession *×* treatment (+Nitrogen/*Rhizobium*).

### Effects of inoculation on plant growth and N accumulation

*Rhizobium* inoculation (R) delayed flowering (DTF) by 2 days on average compared to +N treatment for both cultivated and wild accessions, with a few exceptions (Figure 1A). Cultivated accessions CDC Milestone and ILL 1704 did not flower later when inoculated (R) compared to +N treatment. Wild accessions BGE 016880 (*L. orientalis*), IG 72760 (*L. odemensis*), PI 572376 (*L. orientalis*) and PI 572390 (*L. tomentosus*) also flowered at the same time under both R and +N treatments. IG 72622 (*L. orientalis*) was the only accession that flowered later with the +N treatment compared with the R treatment.

**Figure 1.**
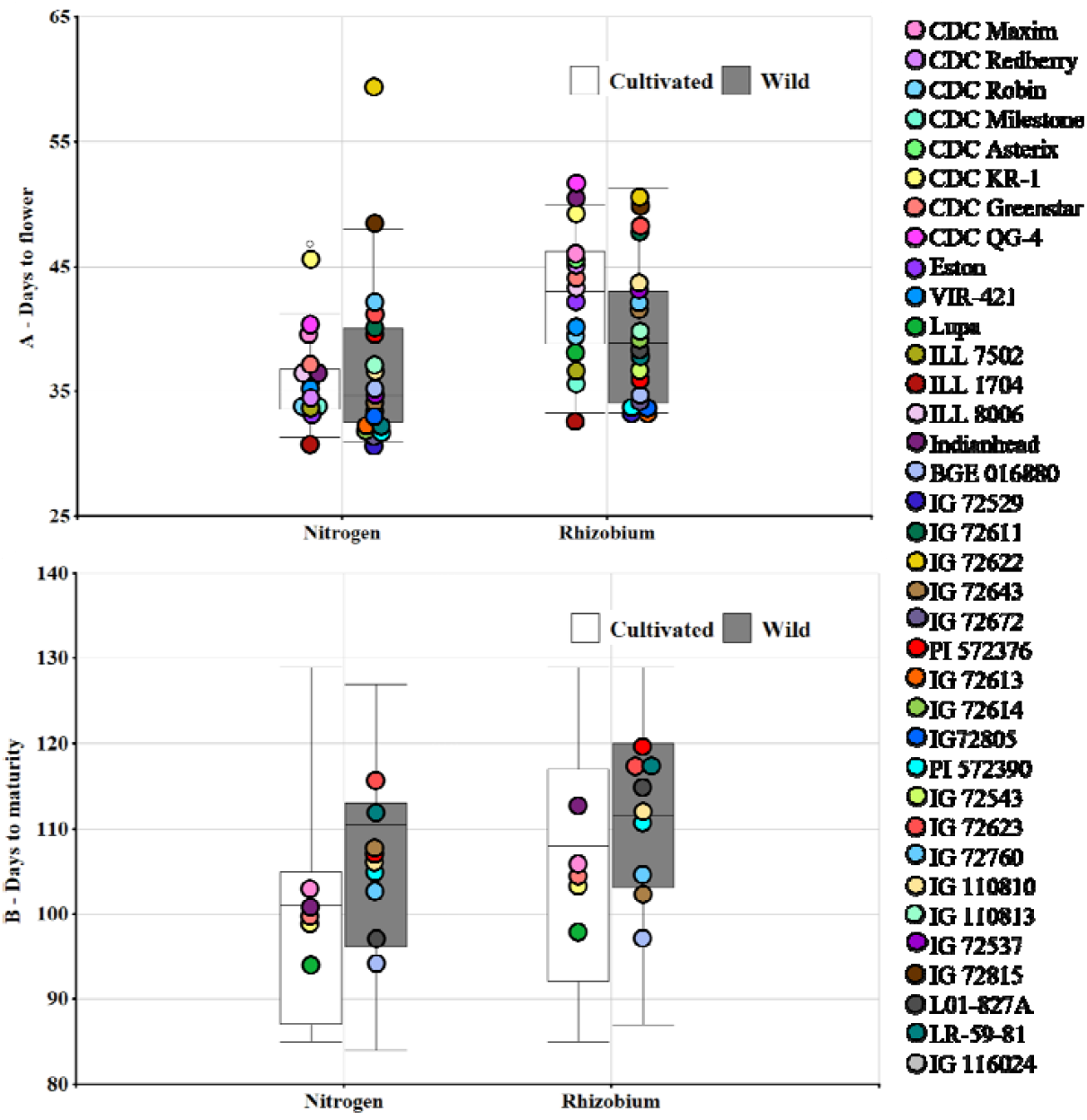
A: days to flowering (LSD 4.8, P=0.01) of 36 *Lens* species accessions, and B: days to maturity (LSD 9.6, P=0.01) of 14 *Lens* species accessions, with added Nitrogen or inoculated with BASF 1435 Nodulator XL^®^ (*Rhizobium leguminosarum* bv. *viciae*) as treatments. Means of cultivated accessions (white boxplots) are displayed separately from the wild ones (gray boxplots).

For the sub-group of 14 accessions studied until maturity (indicated in Table 1), the average days to maturity (DTM) were not different for R and +N treatments (Figure 1B). Among these genotypes, only Indianhead and L01-827A (*L. ervoides*) experienced delays of 11.7 and 18.2 days, respectively, when inoculated with *Rhizobium* (See Supplemental Table S2 for all DTF and DTM means).

Mean shoot dry weight (SDW) was the highest under the +N treatment for both cultivated and wilds (Figure 2A). The R treatment mean was significantly lower and fewer differences were observed among the wild accessions compared to the differences found among cultivated accessions. Plants grown under the +N treatment were generally larger at flowering as measured by shoot and root dry weights (SDW, RDW, Figure 2A. Supplemental Table S3). Some cultivated lines, however, had similar SDW under both +N and R treatments and a couple had similar RDW. Wild accessions were of similar size whether inoculated with *Rhizobium* or given no N (Control treatment). A few cultivated lines had RDW values that were the same whether inoculated or given no N (Control) (Figure 2B, Supplemental TableS3).

**Figure 2.**
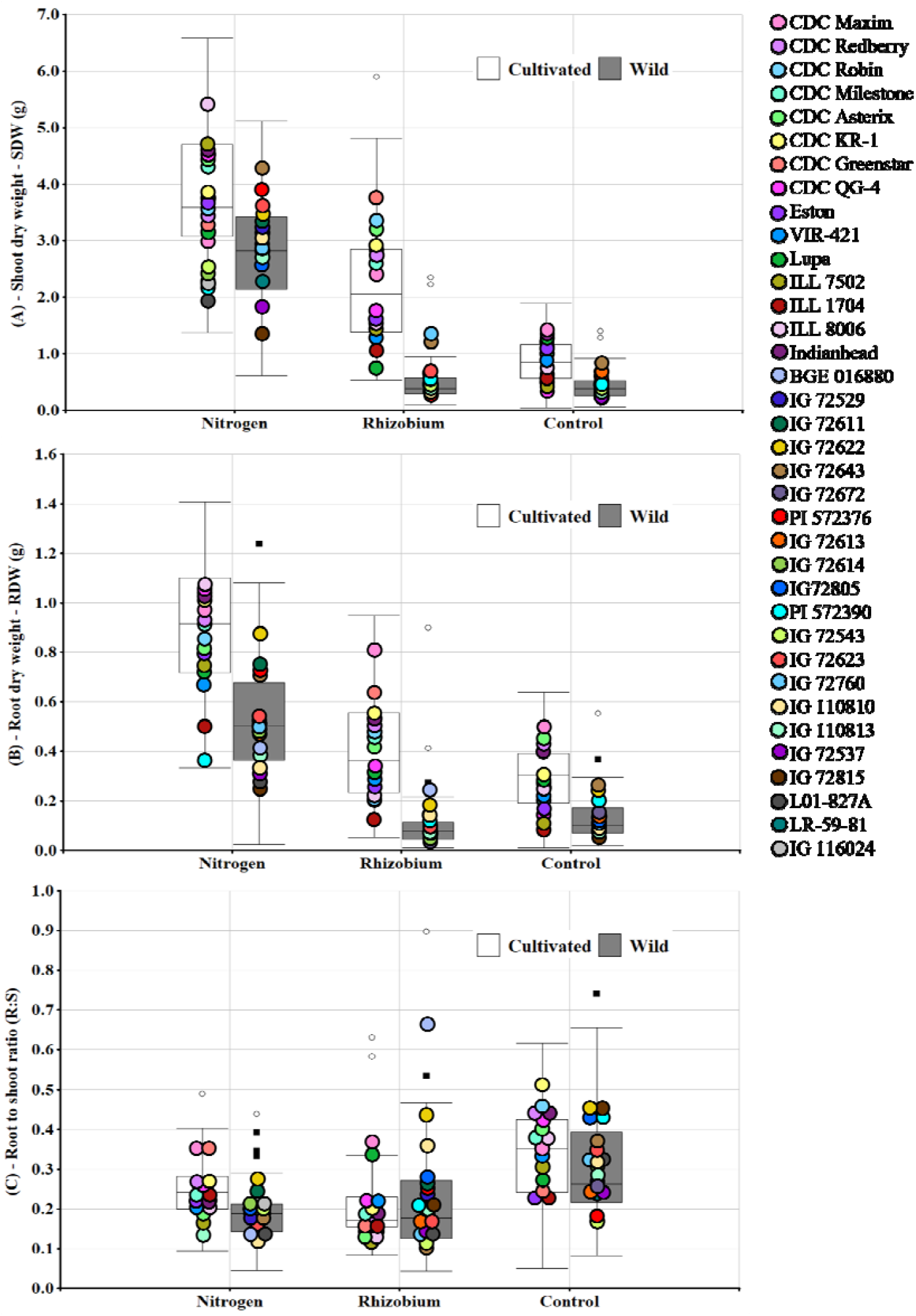
A: shoot dry weight (LSD 0.96, P=0.01), B: root dry weight (LSD 0.21, P=0.01) and C: root to shoot ratio (LSD 0.2, P=0.01), of 36 cultivated and wild *Lens* species accessions with added Nitrogen or inoculated with BASF 1435 Nodulator XL^®^ (*Rhizobium leguminosarum* bv. *viciae*) and Control treatments measured at flowering. Means of cultivated accessions (white boxplots) are displayed separately from the wild ones (gray boxplots).

On average, overall root to shoot ratios (R:S) at flowering were no different across treatments (Figure 2C, Supplemental Table S3), with some individual exceptions. For instance, IG 110810 (*L. lamottei*) and BGE 016880 (*L. orientalis*) had higher ratios when inoculated compared to +N. Some accessions shifted more resources into shoots when inoculated relative to the Control resulting in lower R:S. Among the cultivated lines, CDC Redberry, CDC Robin, and CDC KR-1 fall into this group.

Similarly, a group of the wild accessions, including IG 72643 (*L. orientalis*), PI 572390 (*L. tomentosus*), L01-827A (*L. ervoides*) had lower R:S values when inoculated than were observed when no N was added (Control).

When comparing the amount of N in shoots obtained from the media (as measured under the Control treatment) relative to that obtained from fixation by rhizobia, wild accessions tended to be able to scavenge a greater proportion of N from the media than most cultivated lines (Figure 3). CDC Robin and CDC Milestone were among the cultivated accessions with the highest proportion of N obtained from fixation. IG 72529 (*L. orientalis*), IG 72672 (*L. orientalis*) and IG72614 (*L. tomentosus*) were among the wilds with the lowest proportion of N obtained from fixation.

**Figure 3.**
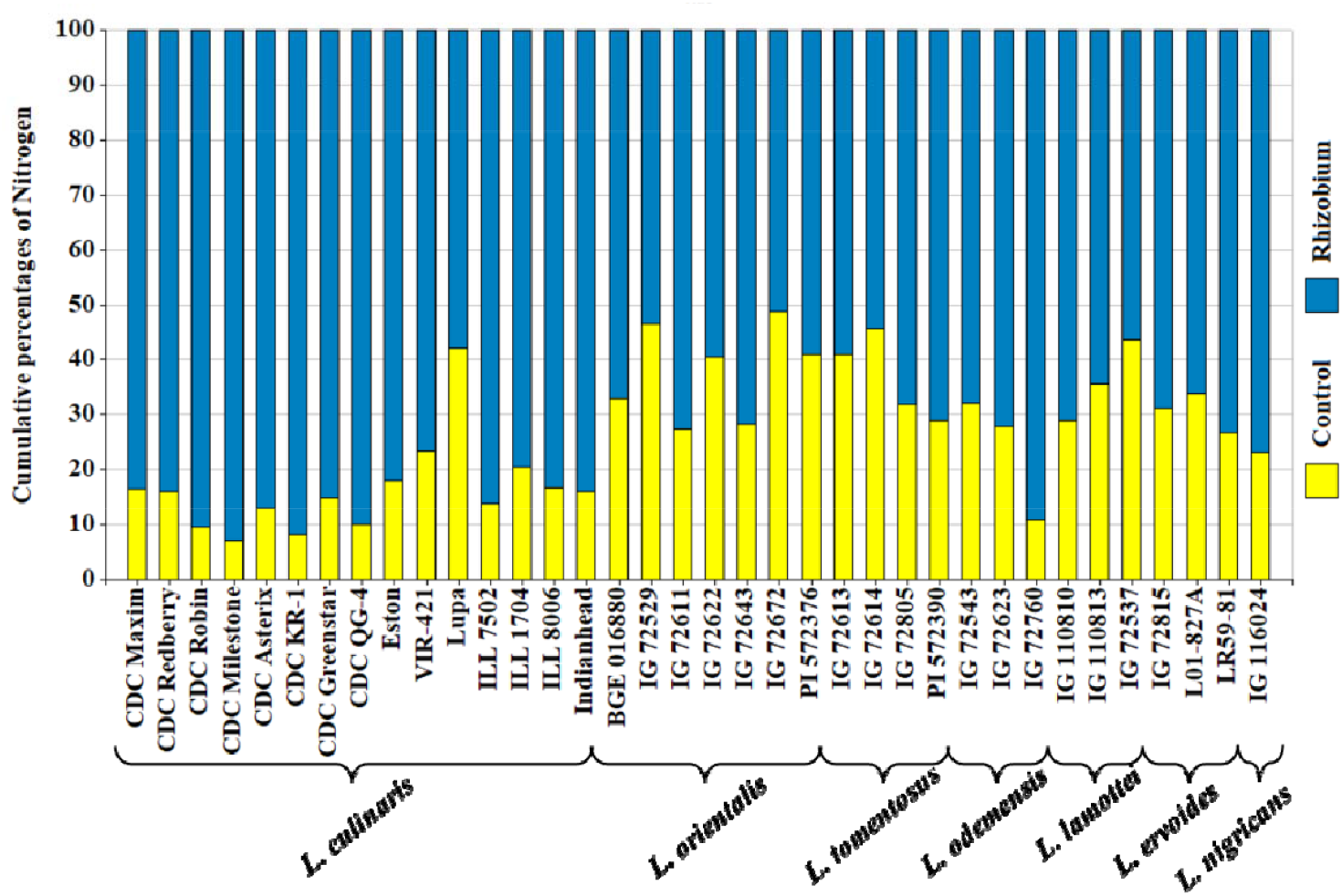
Cumulative percentages of Nitrogen in shoots. Total N accumulation= N_2_ fixed (N yield *Rhizobium* plot-N yield Control plot) + N yield Control plot. N yield Control plot was the mineral N on media.

### Effects of inoculation on root distribution

Most of the accessions responded with higher values for all root parameters with +N treatment. Differences observed between R and C treatments were among the cultivated ones, but none of the wild accessions had differences for any of the parameters under these two treatments.

All the root parameters evaluated were larger under the +N treatment compared to those observed under the R treatment overall and were similar between the R and C treatments (Figure 4, Supplemental Table S4). For the total root length (TRL), CDC Maxim and CDC Robin had similar TRL under both +N and R treatments. Cultivated accessions CDC Maxim, CDC Robin, CDC Milestone, CDC KR-1, CDC Greenstar and Indianhead were the only ones to show differences under the R and C treatments. L01-827A (*L. ervoides*) had similar TRL under +N and C, and IG 72815 (*L. ervoides*) had similar TRL in all treatments.

**Figure 4.**
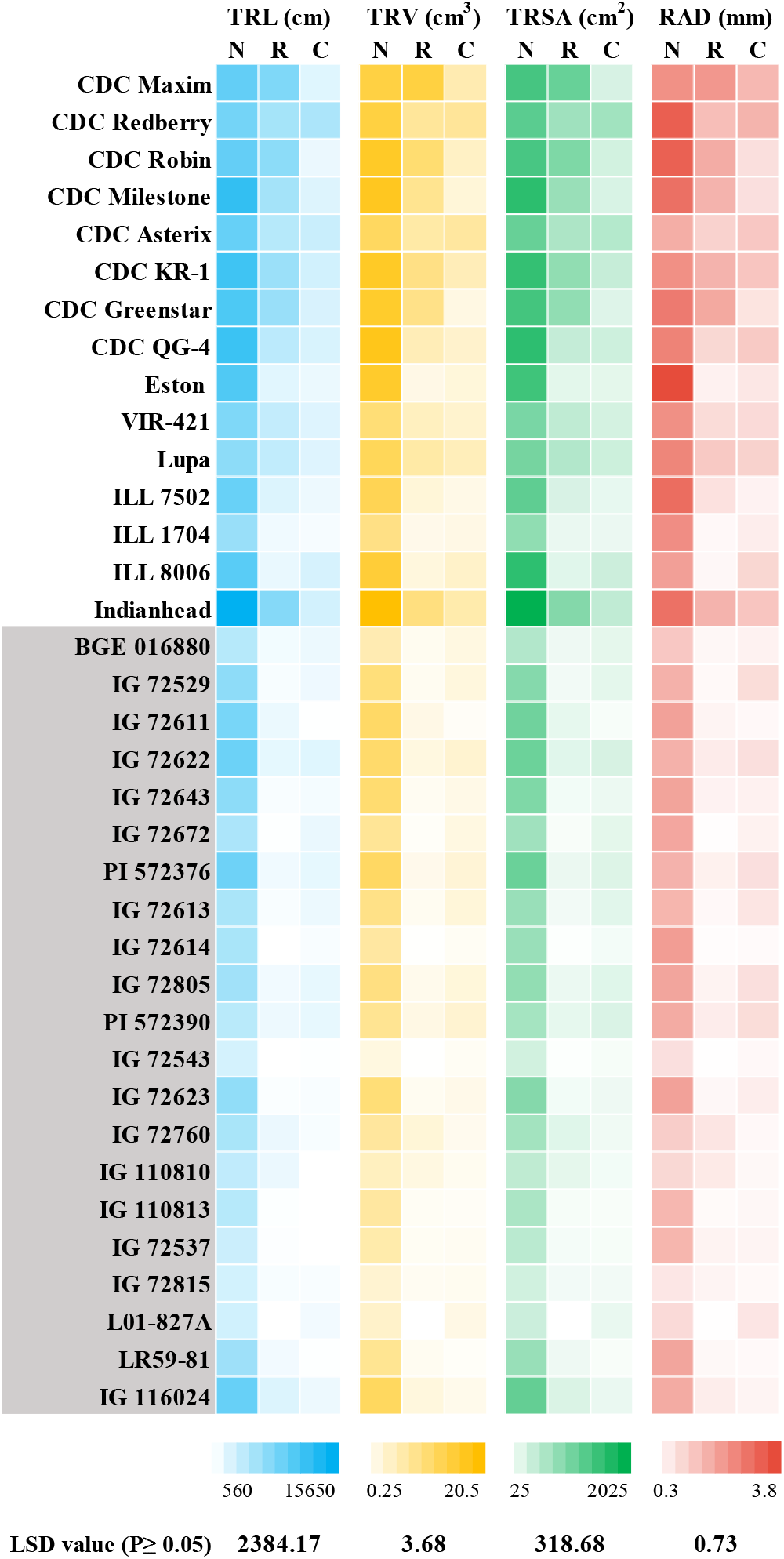
Heatmap for total root length (TRL), total root volume (TRV), total root surface area (TRSA) and root average diameter (RAD) of 36 *Lens* species accessions with added Nitrogen (N), BASF 1435 Nodulator XL^®^ (*Rhizobium leguminosarum* bv. *viciae*) (R) and Control (C) treatments. White: cultivated and grey: wild accessions.

Total root volume (TRV) was different between roots observed with +N and R, except for CDC Maxim, IG 72543 (*L. odemensis*) and IG 110810 (*L. lamottei*) (Figure 4, Supplemental Table S4). Values of TRV were similar under the R and C treatment except for the group of cultivated accessions: CDC Maxim, CDC Robin, CDC Milestone, CDC KR-1 and CDC Greenstar. TRV of L01-827A *(L. ervoides*) and IG 72815 (*L. ervoides*) were not different under any treatment.

Total root surface area (TRSA) observed between +N and R treatments was only similar for CDC Maxim and IG 110810 (*L. lamottei*) and the rest presented significantly lower TRSA under R compared to +N (Figure 4, Supplemental Table S4). Most accessions had similar values under R and C. Only the group of cultivated accessions: CDC Maxim, CDC Robin, CDC Milestone, CDC KR-1, CDC Greenstar and Indianhead, had values of TRSA different between R and C treatments. The TRSA of IG 72543 (*L. odemensis*) was not different under +N and C treatments. IG 72815 (*L. ervoides*) and L01-827A (*L. ervoides*) TRSA did not differ under any treatments.

Root average diameter (RAD) was also similar between +N and R only for a number of accessions: CDC Maxim, CDC Asterix, CDC KR-1, IG 72543 (*L. odemensis*) and IG 72760 (*L. odemensis*) (Figure 4, Supplemental Table S4). The rest of accessions had higher RAD under +N compared to R. RAD observed under R and C treatments was only similar for the accessions CDC Robin, CDC Milestone and CDC Greenstar, and the rest of accessions exhibited similar RAD under both treatments. The RAD of both CDC Asterix and IG 72543 (*L. odemensis*) were similar under +N and C treatments. IG 110810 (*L. lamottei*), IG 72815 (*L. ervoides*) and L01-827A (*L. ervoides*) RAD were not different under any treatment.

### Correlations among evaluated parameters

To understand the relationships among parameters evaluated at flowering and maturity, correlation coefficients based on phenotypic data from the 14 accessions evaluated up to maturity were explored (Figure 5). Correlations among the 36 accessions evaluated at flowering showed similar results among traits evaluated at flowering (Supplemental Figure S1). Nodulation parameters NN, NDW and N fixed, were correlated to KSW. N fixed had the strongest correlation with SDW and NDW (0.95 and 0.72 respectively). SW was correlated with SDW, NDW and N fixed. R:S was only correlated with RDW. N concentration in shoots, SN and seed protein percentages were not correlated with any other measured parameter.

**Figure 5.**
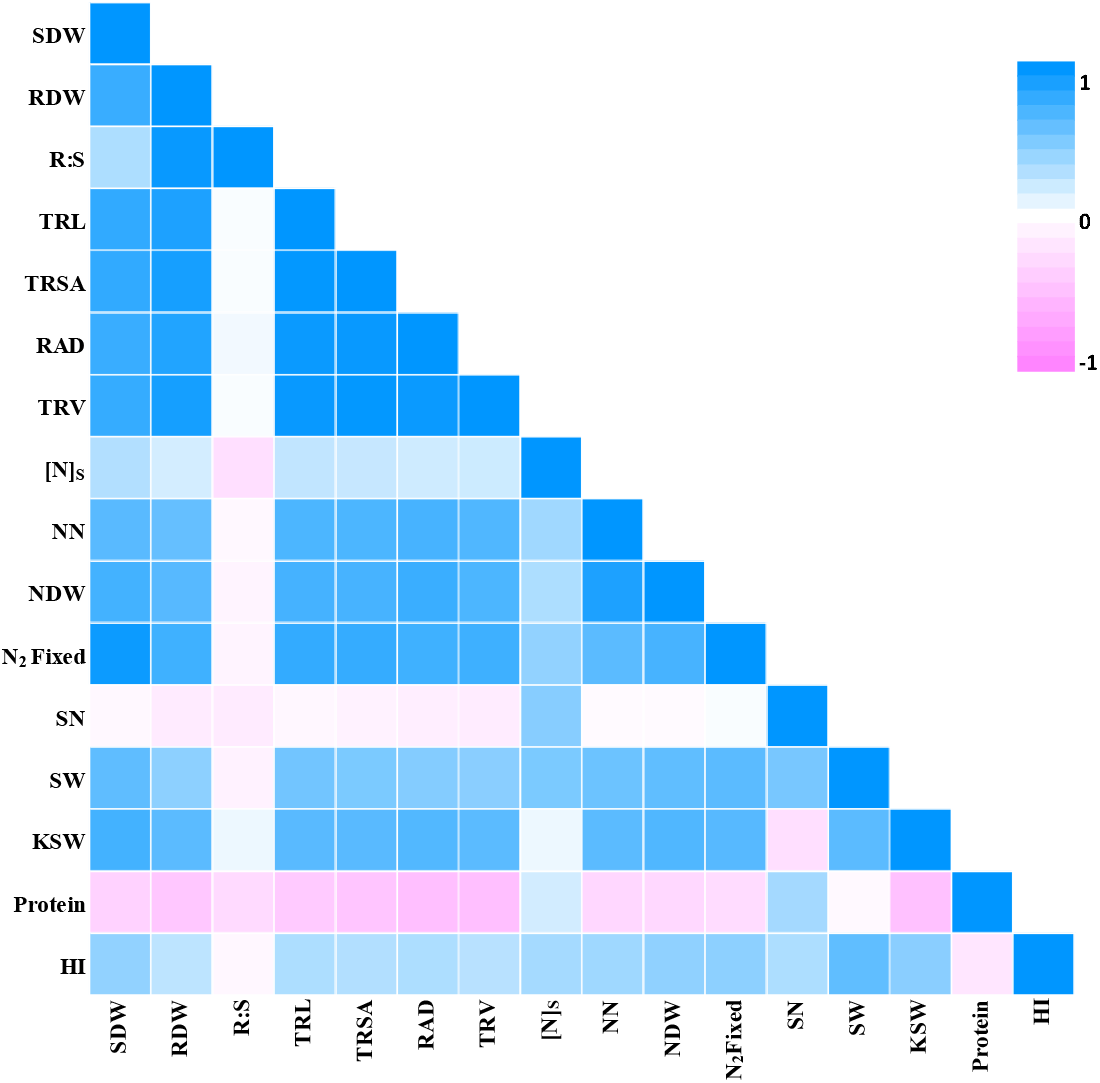
Pearson correlation coefficients among parameters evaluated on 14 *Lens* accessions at flowering and maturity, when inoculated with BASF 1435 Nodulator XL^®^ (*Rhizobium leguminosarum* bv. *viciae*). SDW: shoot dry weight, RDW: root dry weight, R:S: root to shoot ratio, TRL: total root length, TRSA: total root surface area, RAD: root average diameter, TRV: total root volume, Ns: N concentration in shoots, NN: number of nodules, NDW: nodule dry weight, N_2_ fixed, SN: seed number, SW: seed weight, KSW: thousand seed weight, Protein: seed protein percentages, and HI: harvest index.

Shoot dry weight was highly correlated with RDW, all root parameters measured with WinRhizo (TRL, TRSA, RAD and TRV), NDW and N fixed. RDW had the strongest correlation with the root parameters, NDW and N fixed. Root parameters were highly correlated with NN, NDW, N fixed and KSW.

## DISCUSSION

Exploring a greater diversity of *Lens* is necessary to identify new resources within the genus *Lens* to improve the N fixing ability of lentil. The selected group of cultivated accessions are representative of the genetic material generated by the Crop Development Centre (CDC) at the University of Saskatchewan and other accessions representing other growing areas of the world. We also included wild accessions from six related wild species that are already part of the CDC breeding program or are being characterized for other useful traits such as disease resistance. The purpose of the study was to identify any species or specific accessions with characteristics related to higher symbiotic ability.

Both cultivated and wilds displayed broad differences and trends did not correspond to any specific *Lens* species, but this group exhibited some of the differential mechanisms that exist between wild and cultivated plants in their N fixing process and how this impacts seed production. Major dissimilarities were found between wilds and cultivars for phenotypic nodulation system, amount of N fixed and translocated, protein content and seed yield. Differences in root adaptation mechanisms in inoculated plants were also observed.

All vegetative measurements, as well as N fixed, were taken at Reproductive stage 1 (R1) just as flowering started. From R1 to R3, more flowers and vegetative structures continue to develop. CDC Greenstar and Indianhead were among the highest N fixers at flowering with values of fixation above the average for all cultivated accessions. Furthermore, they also had similar seed weight, thousand seed weight and harvest index when inoculated compared to optimum fertilized plants (+N). Conversely, CDC KR-1 was one of the highest N fixers at flowering but yielded the same as CDC Maxim and Lupa, which were intermediate and poor N fixers, respectively. In contrast, wild accessions kept similar seed weights regardless of the amount of N fixed at flowering, except for PI 572376 (*L. orientalis*). The consistent yields did not have any negative effects on seed size, protein content nor harvest index, rather some higher protein content and harvest index were observed. IG 72623 (*L. odemensis*) in particular, fixed 12.4 mg of N at flowering and CDC Maxim fixed 62.3 mg, but both yielded the same. Results of N accumulation in this study suggest that wild *Lens* species have a higher ability to efficiently use N. Studies conducted with 14 cultivated crop species show that only four cereals have higher ability to use N compared to their wild relatives, and most cultivated crops have not outdone their wild relatives (Rotundo and Cipriotti 2016).

For the accessions evaluated up to maturity (R8), the Pearson correlation coefficient between N fixed and seed weight (SW) was 0.64 (Figure 5), a moderate relationship between the two variables. In fact, this was the highest correlation observed between SW and any other parameter measured at flowering, suggesting that selection of genotypes for better N fixing ability cannot be based solely on their phenotypic expression at flowering. Ultimately, what determines a genotype’s superior ability to assimilate N is seed production (Unkovich *et al*., 2008). In addition to this, the higher HI (Table 3) identified among wild accessions (from 4 species) suggests they are more efficient at translocating N to seed. Wild accessions also had overall higher protein concentration in seeds, and at the same time, their protein values were increased when inoculated with *Rhizobium*. Transfer of a higher ability to obtain and translocate N from wild legume species was already known. Successful increases in seed protein content of about 15 % have been obtained in pigeon pea (*Cajanus cajan*) through introgression from wild relatives (Saxena *et al*., 2002). The mechanisms through which they can translocate more N from symbiosis, compared to when they received added N, has to be further studied in lentil.

N fixed values were estimated based on a comparison between each genotype relative to its own reference plant. It is a reliable technique when conducted under low N controlled conditions. However, genotypes varied in their mechanisms to acquire N, consequently, N fixing and reference plants do not remove identical amounts of N from the media. N fixing plants can obtain their N from inorganic N absorbed through roots (added N), fixed N from their bacterial symbiont, and, under very low N availability, organic sources may be accessed from the soil (Lipson and Näsholm 2001). In this experiment the control treatment (C) was expected to contain small amounts of this organic N. As shown in Figure 3, the percentage of N acquired in the Control treatment, in relation to the percentage acquired from the symbiont, tended to be greater in most of the wilds. One exception was IG 72760 (*L. odemensis*), where the N fixed estimate is greater than all the other wild accessions.

It is important to identify the root adaptation mechanisms specific genotypes utilized under N limiting conditions and how this affects N acquisition. Plants responded to changing N resources by altering allocation biomass patterns. More root biomass allocation when lower water or nutrients are available is one of mechanisms plants employ to tolerate these types of stresses (Ågren and Ingestad 1987). In lentil, genotype-specific mechanisms of root allocation have been observed under drought conditions (Gorim and Vandenberg 2017). The significantly higher R:S ratio of some accessions under the Control treatment (Figure 2C) reflects the differential ability of genotypes to use low organic N sources. Those with higher ratios manage to allocate more root biomass to extract more of the N available in the growing media, as was the case of BGE 016880 (*L. orientalis*), IG 72643 (*L. orientalis*) and L01-827A *(L. ervoides*). Some of the cultivars, including CDC Redberry and CDC KR-1, were also among those with higher R:S ratios. In contrast, CDC Maxim, kept the same R:S across treatments. Most cultivars, however, remove similar and low amounts of N regardless of how they allocated their biomass, which shows their inability to adapt to N deficient stress conditions.

The specific root parameters (Figure 4), showed in more detail some of the architectural modifications in the root systems employed by a few accessions in response to the different treatments. Cultivated accessions had smaller values for all parameters with *Rhizobium* inoculation and as expected, under the Control treatment. In contrast, the wilds did not show differences between the inoculation and Control treatments. While cultivars are larger plants in general, inoculation leads to even larger differences. Efficient N fixation makes allocation of root biomass independent of soil N supply (Markham and Zekveld 2007). The smaller root systems observed with *Rhizobium* inoculation, mainly in the wilds, compared to those observed with +N, suggests that they are more efficient at N fixation. The higher allocation of root biomass in the cultivars, when inoculated compared to +N, could also be a determining factor in the lower values of seed yield observed in cultivars but not in the wilds. Overall, the root parameters changed in proportion to RDW with no particular alteration of any specific root dimension under specific treatments. Of note, however, these differences in RDW among genotypes under the different treatments, are not related to a particular species and only distinctive to *L. culinaris*, suggesting that tolerance to low N has been reduced over the course of selection.

For all phenotypic nodulation parameters, differential phenotypes were observed between and within species. *L. culinaris* had the highest overall net values for NN (Table 2) and total N accumulation (See Supplemental Table S4), with CDC varieties having the greatest values among the cultivars. These accessions have all been selected in fields in the same region where this *Rhizobium* strain has been used for at least two decades. Better symbiotic results have been observed in lentils and other crops when locally adapted strains are used (Kurdali *et al*., 1997,

Ferreira and Hungria 2017). Other *L. culinaris* selected elsewhere, including VIR-421 and the ILL lines, performed poorly with this strain. Lupa, a Spanish variety, was particularly inefficient and was the only cultivated accession with undesirable nodulation characteristics in general. IG 72623 (*L. odemensis*) and IG 72760 (*L. odemensis*) had nodulation characteristics similar to those observed in the cultivated lines for both numbers and phenotypic characteristics. However, the other wilds exhibited a range of phenotypic nodule diversity not observed in the cultivars, which was particularly reflected in higher specific nodule dry weight (SpNDW) values in some accessions (Table 2). Those with higher SpNDW had bigger, indeterminate nodules. The wilds with lower SpNDW had both indeterminate and determinate nodules, which made their SpNDW values not different from those observed among the cultivated accessions. Determinate vs. indeterminate nodules have some fundamental structural differences that impact the bacteroid population. Determinate nodules consist of a homogeneous bacteroid population that synchronously senesces (Rolfe and Gresshoff 1988), while indeterminate ones allow the bacterial population to be established in different longitudinal zones, having simultaneously senescent, fixing and a meristematic developing zone (Vassey *et al*., 1990). Such diversity in nodule phenotypes as well as their role on N fixation should be further explored.

In terms of the phenotypic expression of the accessions at flowering, it was expected that wilds would have lower values of SDW compared to most of the cultivated accessions as they are inherently smaller plants. The high response in SDW to +N including the wild accessions was also expected, as it is the easiest route for a plant to acquire N; in contrast to the energy costly process of fixing atmospheric N_2_ (Postgate 1982). For the wild accessions, differences in SDW between inoculated and +N treatments were greater than those observed among the cultivated lines with the same treatments (Figure 2A). At flowering, all wild accessions tended to have a smaller biomass under the *Rhizobium* treatment, with values that were not different from the Control, but they also had the highest values of N concentration (Table S4) in shoots when inoculated, supporting the hypothesis that there is a differential mechanism for N fixation between cultivated and wild accessions.

Considering the cost that establishing a symbiotic relationship may represent for the plant (Walley 2013), phenological stages were monitored. With some exceptions in both cultivated and wild accessions, inoculation compared to +N caused a delay in flowering (Figure 1A, Supplemental Table S1). Ten out of the 36 accessions studied were not different in flowering time between the two treatments. In addition, for the sub-group studied up to maturity, there was no difference in DTM between R and +N treatments for most accessions. Only Indianhead and L01-827A had a delay in DTM, of 11.7 and 18.2 days, respectively, and both also experienced a flowering delay. This could occur if breeding for better BNF, especially in short season areas, therefore, should be taken into consideration and the effects of this symbiotic relationship in the phenological cycle should be determined under field conditions.

Given the level of specificity and differential interactions that have been demonstrated to occur between *Rhizobium* spp. and various legumes (Gunnabo *et al*., 2019; Laguerre, *et al*., 2003; Reyes and Planchon 1995), including cultivated lentil (Abi-Ghanem *et al*., 2011), it would be prudent to assess the ability of these *Lens* species with a greater set of strains. Lentil-nodulating *Rhizobium* from all the agroecosystems where the crop is grown have been characterized (Gai *et al*., unpublished data, 2022; Harun-or Rashid *et al*., 2014) and correspond to several *Rhizobium* species. These isolates will also correspond to the centre of origin of lentil and may help confirm the level of specificity of this diverse set to fix N, especially the wilds.

*Lens* genotypes, from various species, were identified with higher ability to fix N and promising results of seed yield and seed quality traits that can be useful in breeding for better BNF ability in cultivated lentil. CDC Greenstar was identified as the best resources among the cultivated lines for its ability to fix high amounts of N and yield similar amounts of seeds when inoculated as when optimal N was added, with no repercussion in seed quality, nor alterations in days to maturity. IG 72643 (*L. orientalis*) and IG 72623 (*L. odemensis*) represent potential resources for increasing N fixing ability, based on seed yield produced with *Rhizobium* inoculation, but also for their unique mechanisms of N use efficiency. Although good resources were identified within the cultivated group, the unique characteristics found in wild accessions suggest that they can play a role in the improvement of the BNF, contributing to +N higher protein content and higher N use efficiency in cultivated lentil.

## Abbreviations

BNF: biological nitrogen fixation
C: Control
DTF: days to flowering
DTM: days to maturity
HI: harvest index
N: Nitrogen
+N: added Nitrogen
KSW: thousand seed weight
NDW: nodule dry weight
NN: number of nodules
R: *Rhizobium*
RAD: root average diameter
RDW: root dry weight
R:S: root to shoot ratio
SDW: shoot dry weight
SN: seed number
SW: seed weight
SpNDW: specific nodule dry weight
TRL: total root length
TRSA: total root superficial area
TRV: total root volume.

## Supplemental Material Available

Supplemental material for this article is available online.

## Acknowledgments

The research was conducted as part of the Application of Genomics to Innovation in the Lentil Economy (AGILE) project, a Genome Prairie managed project funded by Genome Canada, Western Grains Research Foundation, the Province of Saskatchewan, and the Saskatchewan Pulse Growers. It was also funded in part by an NSERC Industrial Research Chair Award in lentil breeding to Albert Vandenberg. Thanks to Albert Vandenberg for sharing his lentil knowledge through invaluable conversations.

