## Supplementary material for "INCREASING DIVERSITY AMONG *LENS* SPECIES FOR IMPROVING BIOLOGICAL NITROGEN FIXATION IN LENTIL": Supplemental Figure S1.docx

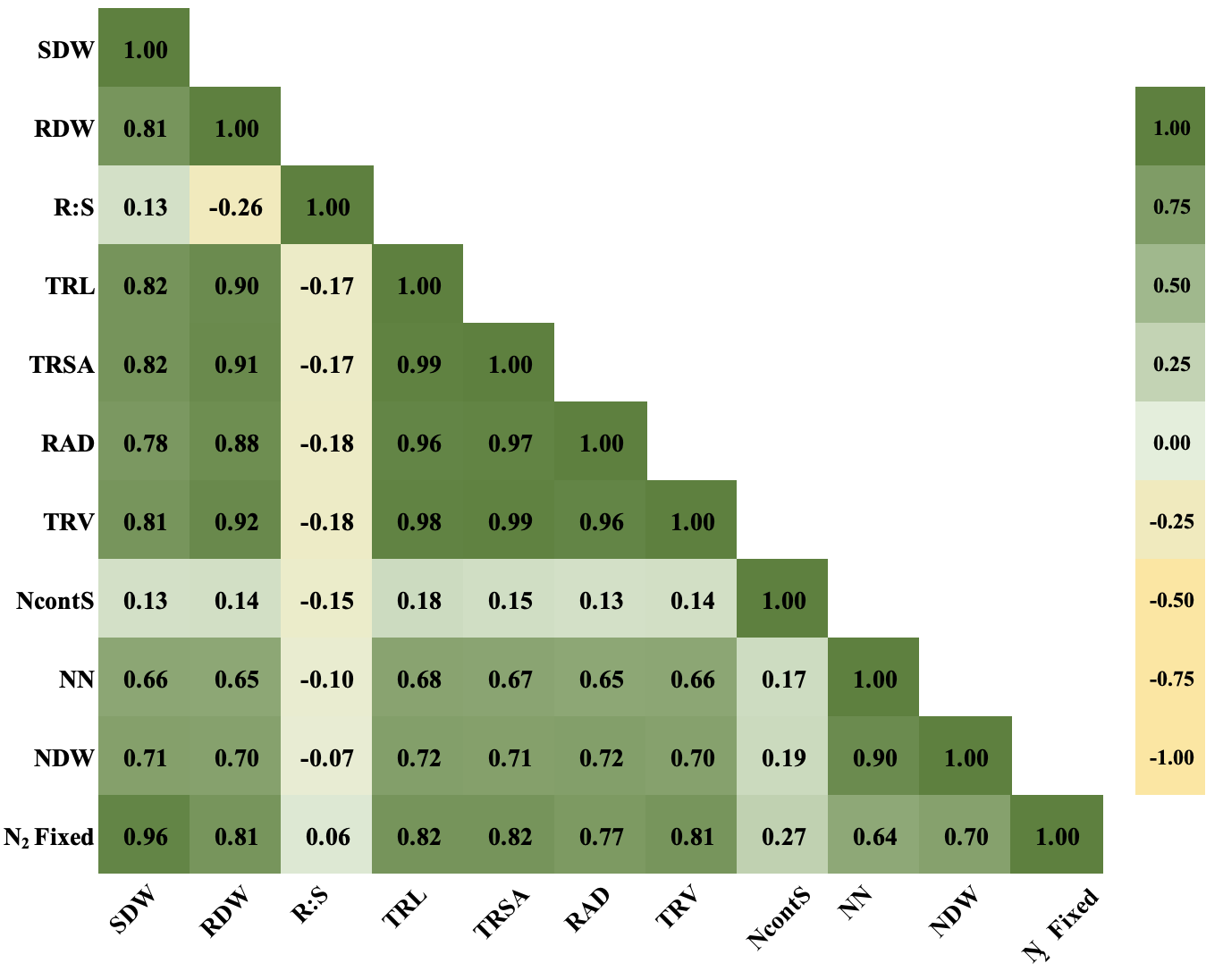

Figure S1. Pearson Correlation among parameters evaluated at flowering of 36 *Lens* species accessions inoculated with BASF 1435 Nodulator XL^®^ (*Rlv*)*.* SDW: *s*hoot dry weight, RDW: root dry weight, R:S: root to shoot ratio, TRL: total root length, TRSA: total root superficial area, RAD: root average diameter, TRV: total root volume, NcontS: N concentration in shoots, NN: number of nodules, NDW: nodule dry weight and N_2_ fixed.
