## Supplementary material for "INCREASING DIVERSITY AMONG *LENS* SPECIES FOR IMPROVING BIOLOGICAL NITROGEN FIXATION IN LENTIL": Supplemental Table S1.docx

| Table S1. Analysis of variance (ANOVA), coefficient of variation (CV) and least significant difference (LSD) values for all evaluated parameters of 36 *Lens* species accessions under added N, BASF 1435 Nodulator XL^®^ (*Rlv*) and control treatments. | | | | | | | |
| --- | --- | --- | --- | --- | --- | --- | --- |
| **ANOVA** | **DTF** | **DTM** | **SDW** | **RDW** | **R:S** | **N_S_** | **N ac** |
| Plot (+N,R,C) | 1.94^***^ | 6.33^***^ | 340.4^***^ | 63.8^***^ | 0.05^**^ | 0.18^***^ | 4.26^***^ |
| Block | 0.94^**^ | ^ns^ | ^ns^ | ^ns^ | ^ns^ | 0.10^***^ | ^ns^ |
| Accession | 2.78^***^ | 5.55^***^ | 556.9^***^ | 121.7^***^ | 0.11^***^ | 0.30^***^ | 14.76^***^ |
| Accession*Plot | 4.82^***^ | 9.61^***^ | 964.5^***^ | 210.7^***^ | 0.2^***^ | 0.52^***^ | 25.56^***^ |
| **ANOVA** | **N_2_ fixed** | **NN** | **NDW** | **SpNDW** | **TRL** | **TRV** | **TRSA** |
| Plot (+N,R,C) |  |  |  |  | 1116.55^***^ | 1.36^***^ | 138.55^***^ |
| Block | 9.84^*^ | ^ns^ | ^ns^ | ^ns^ | ^ns^ | ^ns^ | ^ns^ |
| Accession | 29.52^***^ | 114.2^***^ | 29.72^***^ | 0.28^***^ | 1376.5^***^ | 2.12^***^ | 183.99^***^ |
| Accession*Plot |  |  |  |  | 2384.17^***^ | 3.68^***^ | 318.68^***^ |
| **ANOVA** | **RAD** | **SN** | **SW** | **KSW** | **HI** | **Protein** |  |
| Plot (+N,R,C) | 0.29^***^ | 24.34^***^ | 0.41^***^ | 0.99^***^ | 1.66^***^ | ^ns^ |  |
| Block | ^ns^ | ^ns^ | ^ns^ | ^ns^ | ^ns^ | ^ns^ |  |
| Accession | 0.42^***^ | 46.92^***^ | 0.76^***^ | 2.56^***^ | 4.36^***^ | 0.52^***^ |  |
| Accession*Plot | 0.73^***^ | 81.26^***^ | 1.31^***^ | 4.43^***^ | 7.55^***^ | 0.73^***^ |  |
| ANOVA values are ns: not significant, ^*^, ^**^, ^***^: significant at 0.05, 0.01, 0.001 respectively. DTF: days to flower, DTM: days to maturity, SDW: shoot dry weight, RDW: root dry weight, R:S: root to shoot ratio, N_S_: N concentration in shoots, N ac: N accumulation per plant, NN: number of nodules, NDW: nodule dry weight, SpNDW: specific nodule dry weight, TRL: total root length, TRV: total root volume, TRSA: total root superficial area, RAD: root average diameter, SN: seed number, SW: seed weight, KSW: thousand seed weight, HI: harvest index and seed percentage protein. | | | | | | | |
