## Supplementary material for "INCREASING DIVERSITY AMONG *LENS* SPECIES FOR IMPROVING BIOLOGICAL NITROGEN FIXATION IN LENTIL": Supplemental Table S2.docx

| Table S2. Days to flower and days to maturity of 36 *Lens* species accessions under added Nitrogen, BASF 1435 Nodulator XL^®^ (*Rlv*) and control treatments. | | | | | | |
| --- | --- | --- | --- | --- | --- | --- |
| **Accessions** | **Days to flower** | | | **Days to maturity** | | |
|  | **N** | **R** | **C** | **N** | **R** | **C** |
| CDC Maxim | 40 | 46.3 | 39.3 | 103.17 | 106.2 | 85.3 |
| CDC Redberry | 34 | 44.8 | 40.8 | - | - | - |
| CDC Robin | 33.5 | 39.3 | 38.8 | - | - | - |
| CDC Milestone | 33.5 | 35.3 | 35.5 | - | - | - |
| CDC Asterix | 35.8 | 45.5 | 45 | - | - | - |
| CDC KR-1 | 45.5 | 49 | 44.5 | 99.17 | 103.7 | 88.3 |
| CDC Greenstar | 36.8 | 44.3 | 36 | 99.5 | 104.7 | 85.7 |
| CDC QG-4 | 41.3 | 47.5 | 35.5 | - | - | - |
| Eston | 33 | 42 | 38 | - | - | - |
| VIR-421 | 35 | 40.3 | 36 | - | - | - |
| Lupa | 34.3 | 38.8 | 38.3 | 93.5 | 98.33 | 93.7 |
| ILL 7502 | 33.5 | 36.8 | 34.3 | - | - | - |
| ILL 1704 | 31.3 | 33.3 | 31.8 | - | - | - |
| ILL 8006 | 36.3 | 43 | 41 | - | - | - |
| Indianhead | 36.3 | 50 | 44.5 | 101.5 | 113.2 | 93.7 |
| BGE 016880 | 35.3 | 34.5 | 32.5 | 94.8 | 97 | 87.2 |
| IG 72529 | 31 | 33.3 | 34 | - | - | - |
| IG 72611 | 40.3 | 47.3 | 42.5 | - | - | - |
| IG 72622 | 59 | 51.3 | 48 | - | - | - |
| IG 72643 | 34 | 42.3 | 31.8 | 107.8 | 102.33 | 82.3 |
| IG 72672 | 31.5 | 34 | 33.5 | - | - | - |
| PI 572376 | 40 | 35.8 | 42.5 | 107.7 | 119.7 | 88 |
| IG 72613 | 32.5 | 33.3 | 37 | - | - | - |
| IG 72614 | 32.3 | 39.3 | 37.5 | - | - | - |
| IG 72805 | 33 | 33.5 | 35.8 | - | - | - |
| PI 572390 | 34.3 | 33.5 | 32.5 | 106 | 110.5 | 101.33 |
| IG 72543 | 33.3 | 36.3 | 34.5 | - | - | - |
| IG 72623 | 41.8 | 47.5 | 49 | 115.33 | 118 | 92.7 |
| IG 72760 | 42.3 | 42 | 47 | 101.83 | 103.7 | 88.3 |
| IG 110810 | 36.3 | 44 | 35.8 | 106.7 | 112.3 | 97.2 |
| IG 110813 | 36.5 | 39.5 | 39.8 | - | - | - |
| IG 72537 | 35 | 43.8 | 36 | - | - | - |
| IG 72815 | 48 | 51 | 43.8 | - | - | - |
| L01-827A | 33.5 | 38.3 | 43.5 | 97.8 | 116 | 85.7 |
| LR59-81 | 32.5 | 38.5 | 35.3 | 111.8 | 118 | 113 |
| IG 116024^‡^ | - | - | - | - | - | - |
| **Mean** | 38.6 | 41 | 36.6 | 103.3 | 108.8 | 91.6 |
| LSD | 4.81^***^ | | | 9.61^***^ | | |
| ^‡^IG 116024 did not flower during the time of the experiment.  ANOVA values are ns: not significant, ^*^, ^**^, ^***^: significant at 0.05, 0.01, 0.001 respectively. | | | | | | |
