## Supplementary material for "INCREASING DIVERSITY AMONG *LENS* SPECIES FOR IMPROVING BIOLOGICAL NITROGEN FIXATION IN LENTIL": Supplemental Table S3.docx

| Table S3. Shoot dry weight (mg), root dry weight(mg) and Root:shoot ratio of 36 *Lens* species accessions under added Nitrogen, BASF 1435 Nodulator XL^®^ (*Rlv*) and control treatments. | | | | | | | | | |
| --- | --- | --- | --- | --- | --- | --- | --- | --- | --- |
| **Accessions** | **Shoot dry weight (mg)** | | | **Root dry weight (mg)** | | | **Root:shoot ratio** | | |
|  | **N** | **R** | **C** | **N** | **R** | **C** | **N** | **R** | **C** |
| CDC Maxim | 2980.0 | 2370.0 | 1347.5 | 995.6 | 807.7 | 487.5 | 0.35 | 0.38 | 0.35 |
| CDC Redberry | 3455.0 | 2827.5 | 1082.5 | 958.6 | 506.6 | 433.9 | 0.27 | 0.18 | 0.44 |
| CDC Robin | 3515.0 | 3420.0 | 652.5 | 857.5 | 501.2 | 263.3 | 0.25 | 0.15 | 0.47 |
| CDC Milestone | 4250.0 | 2760.0 | 532.5 | 951.6 | 493.9 | 205.9 | 0.24 | 0.19 | 0.38 |
| CDC Asterix | 4475.0 | 3110.0 | 1087.5 | 820.0 | 440.3 | 435.2 | 0.19 | 0.14 | 0.40 |
| CDC KR-1 | 3827.5 | 2930.0 | 687.5 | 1027.4 | 570.6 | 343.1 | 0.27 | 0.20 | 0.51 |
| CDC Greenstar | 3200.0 | 3837.5 | 1317.5 | 1094.3 | 623.8 | 276.3 | 0.35 | 0.16 | 0.23 |
| CDC QG-4 | 4602.5 | 1892.5 | 420.0 | 1102.4 | 377.1 | 179.8 | 0.26 | 0.22 | 0.43 |
| Eston | 3595.0 | 1717.5 | 942.5 | 814.6 | 258.7 | 181.4 | 0.23 | 0.15 | 0.22 |
| VIR-421 | 3787.5 | 1285.0 | 775.0 | 647.1 | 291.4 | 239.2 | 0.20 | 0.22 | 0.34 |
| Lupa | 3160.0 | 897.7 | 1227.5 | 695.7 | 335.8 | 321.2 | 0.23 | 0.34 | 0.27 |
| ILL 7502 | 4707.5 | 1555.0 | 522.5 | 761.5 | 202.6 | 149.2 | 0.17 | 0.13 | 0.31 |
| ILL 1704 | 2237.5 | 1037.5 | 570.0 | 507.9 | 160.1 | 130.8 | 0.24 | 0.16 | 0.22 |
| ILL 8006 | 5485.0 | 1715.0 | 745.0 | 1104.9 | 220.6 | 274.4 | 0.20 | 0.14 | 0.38 |
| Indianhead | 4660.0 | 2840.0 | 1342.5 | 1087.9 | 567.6 | 427.4 | 0.23 | 0.19 | 0.35 |
| BGEO 16880 | 2910.0 | 480.0 | 480.0 | 412.5 | 293.1 | 163.9 | 0.14 | 0.66 | 0.34 |
| IG 72529 | 3210.0 | 362.5 | 587.5 | 535.3 | 79.2 | 153.0 | 0.17 | 0.22 | 0.28 |
| IG 72611 | 3312.5 | 557.5 | 210.0 | 771.3 | 136.7 | 71.9 | 0.25 | 0.28 | 0.33 |
| IG 72622 | 3462.5 | 455.0 | 490.0 | 888.9 | 194.9 | 219.7 | 0.29 | 0.43 | 0.45 |
| IG 72643 | 4267.5 | 1127.5 | 902.5 | 706.6 | 88.5 | 237.7 | 0.17 | 0.10 | 0.38 |
| IG 72672 | 2855.0 | 432.5 | 727.5 | 517.4 | 61.3 | 174.9 | 0.18 | 0.16 | 0.27 |
| PI 572376 | 3955.0 | 350.0 | 517.5 | 731.5 | 97.7 | 105.3 | 0.19 | 0.27 | 0.19 |
| IG 72613 | 3215.0 | 447.5 | 765.0 | 503.2 | 75.4 | 173.3 | 0.18 | 0.17 | 0.24 |
| IG 72614 | 2360.0 | 152.5 | 225.0 | 471.5 | 27.7 | 68.2 | 0.21 | 0.19 | 0.31 |
| IG 72805 | 2547.5 | 472.5 | 367.5 | 508.4 | 86.6 | 153.3 | 0.20 | 0.29 | 0.43 |
| PI 572390 | 2122.5 | 572.5 | 565.0 | 386.4 | 124.3 | 210.0 | 0.18 | 0.21 | 0.43 |
| IG 72543 | 2557.5 | 475.0 | 462.5 | 537.1 | 52.0 | 83.4 | 0.20 | 0.11 | 0.18 |
| IG 72623 | 3697.5 | 607.5 | 422.5 | 575.7 | 90.6 | 138.4 | 0.16 | 0.17 | 0.36 |
| IG 72760 | 2905.0 | 1340.0 | 330.0 | 487.9 | 204.3 | 102.6 | 0.17 | 0.14 | 0.33 |
| IG 110810 | 3085.0 | 397.5 | 267.5 | 342.5 | 148.8 | 81.9 | 0.12 | 0.36 | 0.32 |
| IG 110813 | 2822.5 | 437.5 | 350.0 | 390.1 | 81.8 | 66.6 | 0.13 | 0.20 | 0.19 |
| IG 72537 | 1882.5 | 310.0 | 292.5 | 347.1 | 41.9 | 71.6 | 0.18 | 0.15 | 0.24 |
| IG 72815 | 1297.8 | 280.0 | 152.5 | 223.1 | 62.3 | 66.4 | 0.17 | 0.21 | 0.45 |
| L01-827A | 1967.5 | 147.5 | 355.0 | 272.1 | 18.2 | 128.3 | 0.14 | 0.14 | 0.34 |
| LR59-81 | 2377.5 | 632.5 | 322.5 | 502.8 | 95.6 | 60.1 | 0.18 | 0.14 | 0.24 |
| IG 116024 | 2205.0 | 460.0 | 240.0 | 741.1 | 157.6 | 100.2 | 0.35 | 0.34 | 0.41 |
| **Mean** |  |  |  | 0.676 | 0.238 | 0.194 | 0.21 | 0.22 | 0.33 |
| LSD | 0.96^***^ | | | 0.21^***^ | | | 0.2^***^ | | |
| ANOVA values are ns: not significant, ^*^, ^**^, ^***^: significant at 0.05, 0.01, 0.001 respectively. | | | | | | | | | |
