## Supplementary material for "INCREASING DIVERSITY AMONG *LENS* SPECIES FOR IMPROVING BIOLOGICAL NITROGEN FIXATION IN LENTIL": Supplemental Table S4.docx

| Table S4. Total root length, total root volume total root superficial area and root average diameter of 36 *Lens* species accessions with added Nitrogen, BASF 1435 Nodulator XL^®^ (*Rlv*) and control treatments. | | | | | | | | | | | | |
| --- | --- | --- | --- | --- | --- | --- | --- | --- | --- | --- | --- | --- |
| **Accessions** | **Total root length (cm)** | | | **Total root volume (cm^3^)** | | | **Total root superficial area (cm^2^)** | | | **Root average diameter (mm)** | | |
|  | **N** | **R** | **C** | **N** | **R** | **C** | **N** | **R** | **C** | **N** | **R** | **C** |
| CDC Maxim | 9847.0 | 8123.0 | 2444.1 | 15.2 | 15.3 | 6.5 | 1476.0 | 1212.6 | 340.5 | 2.5 | 2.3 | 1.7 |
| CDC Redberry | 8730.5 | 5876.4 | 5506.8 | 15.3 | 8.2 | 8.3 | 1311.7 | 778.0 | 755.3 | 3.4 | 1.6 | 1.8 |
| CDC Robin | 9746.5 | 7418.0 | 1719.7 | 17.3 | 11.5 | 4.8 | 1453.7 | 1032.4 | 374.5 | 3.4 | 1.9 | 0.9 |
| CDC Milestone | 12638.9 | 5983.9 | 2615.9 | 17.9 | 9.1 | 3.3 | 1681.8 | 827.2 | 327.2 | 3.1 | 1.8 | 0.9 |
| CDC Asterix | 9570.8 | 4909.2 | 3800.2 | 12.9 | 7.3 | 7.9 | 1216.7 | 666.4 | 607.3 | 1.9 | 1.2 | 1.4 |
| CDC KR-1 | 11940.1 | 6432.6 | 3288.2 | 17.5 | 10.0 | 5.9 | 1615.6 | 895.5 | 471.9 | 2.5 | 1.8 | 1.4 |
| CDC Greenstar | 11056.1 | 6560.5 | 2925.8 | 16.9 | 9.7 | 2.5 | 1494.1 | 891.0 | 287.2 | 2.9 | 2.0 | 0.8 |
| CDC QG-4 | 12106.5 | 4582.4 | 2841.5 | 18.4 | 6.0 | 4.3 | 1670.6 | 489.6 | 414.9 | 2.8 | 1.1 | 1.3 |
| Eston | 10946.8 | 2297.7 | 1698.4 | 17.1 | 2.2 | 3.1 | 1525.7 | 250.7 | 243.5 | 3.8 | 0.6 | 0.8 |
| VIR-421 | 8118.2 | 4159.0 | 2541.2 | 11.2 | 5.3 | 4.1 | 1063.1 | 524.7 | 359.6 | 2.5 | 1.0 | 1.0 |
| Lupa | 7318.2 | 4187.9 | 2567.9 | 13.4 | 7.3 | 5.7 | 1102.6 | 621.7 | 426.4 | 2.7 | 1.4 | 1.2 |
| ILL 7502 | 9480.2 | 2653.3 | 1566.3 | 13.8 | 3.3 | 2.1 | 1272.3 | 329.7 | 206.9 | 3.1 | 0.9 | 0.6 |
| ILL 1704 | 6562.6 | 1465.4 | 1142.3 | 10.0 | 2.0 | 2.4 | 908.5 | 189.8 | 182.5 | 2.5 | 0.4 | 0.6 |
| ILL 8006 | 10414.6 | 1865.2 | 2993.6 | 16.1 | 3.0 | 4.6 | 1667.7 | 263.8 | 430.8 | 2.2 | 0.5 | 1.1 |
| Indianhead | 15644.7 | 7785.1 | 3263.0 | 20.5 | 10.4 | 6.9 | 2021.2 | 1003.4 | 514.5 | 3.1 | 1.8 | 1.4 |
| BGEO 16880 | 4881.2 | 1272.9 | 1653.4 | 6.4 | 1.5 | 2.6 | 623.5 | 155.1 | 232.4 | 1.4 | 0.5 | 0.6 |
| IG72529 | 7249.1 | 1043.4 | 1545.8 | 10.7 | 1.4 | 3.0 | 984.1 | 133.9 | 240.6 | 1.8 | 0.4 | 1.0 |
| IG72611 | 8577.7 | 1699.4 | 655.9 | 12.4 | 2.2 | 1.0 | 1152.0 | 215.9 | 88.8 | 2.1 | 0.5 | 0.4 |
| IG72622 | 9224.4 | 2041.5 | 2411.8 | 12.1 | 2.7 | 4.0 | 1178.8 | 261.3 | 346.3 | 1.8 | 0.7 | 0.9 |
| IG72643 | 7316.6 | 960.6 | 1176.8 | 11.6 | 1.3 | 2.1 | 1028.8 | 123.4 | 173.7 | 2.1 | 0.5 | 0.6 |
| IG72672 | 5470.7 | 730.5 | 1818.9 | 8.7 | 0.8 | 2.6 | 769.1 | 83.6 | 243.8 | 2.0 | 0.3 | 0.6 |
| PI 572376 | 9083.2 | 1506.3 | 2013.7 | 12.7 | 1.9 | 3.5 | 1201.0 | 187.8 | 293.6 | 1.8 | 0.6 | 0.9 |
| IG72613 | 5594.2 | 1007.6 | 1647.9 | 9.8 | 1.4 | 3.2 | 819.4 | 133.9 | 256.8 | 1.7 | 0.4 | 0.8 |
| IG72614 | 5653.8 | 412.8 | 1011.9 | 7.7 | 0.5 | 1.1 | 821.5 | 48.8 | 114.7 | 2.2 | 0.4 | 0.4 |
| Table S4. Continued. | | | | | | | | | | | | |
| IG72805 | 6105.0 | 1397.0 | 1934.3 | 10.3 | 1.8 | 3.1 | 885.9 | 175.0 | 273.3 | 2.1 | 0.5 | 0.9 |
| PI 572390 | 4699.8 | 1571.7 | 1965.9 | 9.0 | 2.4 | 4.0 | 725.0 | 217.6 | 307.9 | 2.0 | 0.7 | 1.0 |
| IG72543 | 3061.4 | 514.0 | 699.7 | 2.7 | 0.4 | 1.1 | 390.7 | 49.2 | 97.7 | 0.9 | 0.3 | 0.4 |
| IG72623 | 7111.6 | 848.8 | 967.1 | 11.1 | 1.4 | 2.1 | 992.8 | 123.2 | 157.5 | 2.1 | 0.5 | 0.6 |
| IG 72760 | 5650.0 | 1785.5 | 1052.7 | 8.1 | 3.5 | 1.6 | 751.0 | 276.4 | 144.4 | 1.3 | 0.8 | 0.4 |
| IG 110810 | 4283.7 | 1732.1 | 574.5 | 5.4 | 2.6 | 1.5 | 533.9 | 236.1 | 121.8 | 1.1 | 0.7 | 0.5 |
| IG 110813 | 4915.6 | 743.3 | 564.9 | 7.8 | 1.1 | 1.0 | 686.2 | 99.1 | 84.3 | 1.7 | 0.4 | 0.5 |
| IG 72537 | 3656.2 | 784.2 | 608.8 | 7.0 | 1.1 | 1.2 | 564.7 | 105.6 | 96.4 | 1.7 | 0.5 | 0.5 |
| IG 72815 | 3206.2 | 1063.0 | 994.8 | 3.9 | 1.2 | 1.5 | 395.7 | 127.8 | 134.0 | 0.8 | 0.5 | 0.4 |
| L01-827A | 3366.7 | 268.7 | 1360.1 | 4.5 | 0.3 | 2.4 | 434.8 | 28.8 | 200.7 | 1.0 | 0.3 | 0.8 |
| LR59-81 | 6340.5 | 1349.5 | 691.4 | 8.8 | 1.4 | 0.9 | 836.2 | 153.0 | 87.5 | 2.1 | 0.5 | 0.4 |
| IG 116024 | 9550.8 | 2640.4 | 1583.3 | 13.0 | 3.0 | 1.8 | 1245.3 | 312.2 | 190.2 | 2.0 | 0.7 | 0.5 |
| **Mean** | 7753.1 | 2768.7 | 1884.7 | 11.6 | 4.0 | 3.3 | 1069.3 | 367.3 | 273.0 | 2.2 | 0.9 | 0.8 |
| LSD _(P≥0.05)_ | 2384.17^***^ | | | 3.68^***^ | | | 318.68^***^ | | | 0.73^***^ | | |
| ANOVA values are ns: not significant, ^*^, ^**^, ^***^: significant at 0.05, 0.01, 0.001 respectively. | | | | | | | | | | | | |
