## Supplementary material for "INCREASING DIVERSITY AMONG *LENS* SPECIES FOR IMPROVING BIOLOGICAL NITROGEN FIXATION IN LENTIL": Supplemental Table S5.docx

| Table S5. N concentration in shoots (%) and total N accumulation (mg/plant) at flowering of 36 *Lens* species accession with added Nitrogen, BASF 1435 Nodulator XL^®^ (*Rlv*) and control treatments. | | | | | | |
| --- | --- | --- | --- | --- | --- | --- |
| **Accessions** | **N concentration (%)** | | | **Total N accumulation (mg/plant)** | | |
|  | **N** | **R** | **C** | **N** | **R** | **C** |
| CDC Maxim | 1.79 | 3.25 | 1.13 | 54.07 | 77.54 | 15.29 |
| CDC Redberry | 2.41 | 3.17 | 1.55 | 81.83 | 89.54 | 17.18 |
| CDC Robin | 1.69 | 2.55 | 1.32 | 59.43 | 86.30 | 9.14 |
| CDC Milestone | 1.98 | 2.95 | 1.16 | 88.35 | 82.55 | 6.23 |
| CDC Asterix | 2.46 | 3.23 | 1.42 | 112.77 | 101.86 | 15.24 |
| CDC KR-1 | 2.42 | 3.47 | 1.36 | 89.79 | 102.37 | 9.23 |
| CDC Greenstar | 1.61 | 2.9 | 1.55 | 50.93 | 110.96 | 19.51 |
| CDC QG-4 | 2.41 | 3.08 | 1.49 | 111.11 | 56.93 | 6.33 |
| Eston | 2.1 | 2.67 | 1.13 | 75.23 | 45.73 | 10.14 |
| VIR-421 | 2.3 | 2.93 | 1.49 | 91.24 | 37.44 | 11.47 |
| Lupa | 1.52 | 1.19 | 1.1 | 45.18 | 13.14 | 9.55 |
| ILL 7502 | 1.47 | 2.2 | 1.04 | 69.24 | 34.46 | 5.48 |
| ILL 1704 | 2.42 | 2.58 | 1.25 | 52.51 | 26.31 | 6.80 |
| ILL 8006 | 2.14 | 2.41 | 1.17 | 117.53 | 41.17 | 8.21 |
| Indianhead | 2.23 | 3.54 | 1.53 | 104.15 | 101.24 | 19.25 |
| BGEO 16880 | 1.83 | 2.22 | 1.1 | 51.91 | 10.67 | 5.22 |
| IG72529 | 1.8 | 3.18 | 1.9 | 54.66 | 11.67 | 10.20 |
| IG72611 | 1.77 | 2.61 | 2.22 | 57.58 | 12.58 | 4.72 |
| IG72622 | 2.03 | 3.16 | 1.97 | 73.09 | 14.40 | 9.86 |
| IG72643 | 2.06 | 2.77 | 1.39 | 86.16 | 30.57 | 12.02 |
| IG72672 | 1.96 | 2.88 | 1.59 | 53.47 | 12.43 | 11.87 |
| PI 572376 | 1.56 | 3.26 | 1.54 | 61.46 | 11.27 | 7.81 |
| IG72613 | 2.23 | 3.4 | 1.42 | 71.51 | 15.16 | 10.51 |
| IG72614 | 2.61 | 2.53 | 1.42 | 62.41 | 3.91 | 3.28 |
| IG72805 | 2.32 | 3.5 | 1.32 | 58.09 | 13.25 | 6.17 |
| PI 572390 | 2.82 | 3.26 | 1.26 | 57.54 | 18.54 | 7.58 |
| IG72543 | 2.67 | 2.5 | 1.19 | 63.73 | 11.96 | 5.65 |
| IG72623 | 2.13 | 3.34 | 1.7 | 78.74 | 20.21 | 7.80 |
| IG 72760 | 2.04 | 3.02 | 1.6 | 58.93 | 40.08 | 4.88 |
| IG 110810 | 1.94 | 2.59 | 1.59 | 57.95 | 10.40 | 4.26 |
| IG 110813 | 1.94 | 2.67 | 1.66 | 54.95 | 10.47 | 5.79 |
| IG 72537 | 2.51 | 2.06 | 1.72 | 45.51 | 6.52 | 5.04 |
| IG 72815 | 2.98 | 2.36 | 1.85 | 38.55 | 6.24 | 2.84 |
| L01-827A | 1.76 | 2.27 | 1.99 | 25.76 | 6.63 | 3.38 |
| LR59-81 | 2.42 | 2.85 | 1.91 | 46.86 | 16.67 | 6.11 |
| IG 116024 | 2.1 | 3.86 | 2.3 | 47.57 | 17.72 | 5.36 |
| **Mean** | 2.12 | 2.84 | 1.51 | 66.94 | 36.17 | 8.78 |
| LSD | 0.52^***^ | | | 25.56^***^ | | |
| ANOVA values are ns: not significant, ^*^, ^**^, ^***^: significant at 0.05, 0.01, 0.001 respectively. | | | | | | |
